# Fighting antimicrobial resistance in *Pseudomonas aeruginosa* with machine learning-enabled molecular diagnostics

**DOI:** 10.1101/643676

**Authors:** Ariane Khaledi, Aaron Weimann, Monika Schniederjans, Ehsaneddin Asgari, Tzu-Hao Kuo, Antonio Oliver, Gabriel Cabot, Axel Kola, Petra Gastmeier, Michael Hogardt, Daniel Jonas, Mohammad R.K. Mofrad, Andreas Bremges, Alice C. McHardy, Susanne Häussler

## Abstract

The growing importance of antibiotic resistance on clinical outcomes and cost of care underscores the need for optimization of current diagnostics. For a number of bacterial species antimicrobial resistance can be unambiguously predicted based on their genome sequence. In this study, we sequenced the genomes and transcriptomes of 414 drug-resistant clinical *Pseudomonas aeruginosa* isolates. By training machine learning classifiers on information about the presence or absence of genes, their sequence variation, and gene expression profiles, we generated predictive models and identified biomarkers of susceptibility or resistance to four commonly administered antimicrobial drugs. Using these data types alone or in combination resulted in high (0.8-0.9) or very high (>0.9) sensitivity and predictive values, where the relative contribution of the different categories of biomarkers strongly depended on the antibiotic. For all drugs except for ciprofloxacin, gene expression information substantially improved diagnostic performance. Our results pave the way for the development of a molecular resistance profiling tool that reliably predicts antimicrobial susceptibility based on genomic and transcriptomic markers. The implementation of a molecular susceptibility test system in routine clinical microbiology diagnostics holds promise to provide earlier and more detailed information on antibiotic resistance profiles of bacterial pathogens and thus could change how physicians treat bacterial infections.

## Introduction

The rise of antibiotic resistance is a public health issue of greatest importance (Cassini et al. 2019). Growing resistance hampers the use of conventional antibiotics and leads to increased rates of ineffective empiric antimicrobial therapy. If not adequately treated, infections cause suffering, incapacity and death, and impose an enormous financial burden on healthcare systems and on society in general (Alanis 2005; Fair and Tor 2014; Gootz 2010). Despite growing medical need, FDA approvals of new antibacterial agents have substantially decreased over the last 20 years (Kinch et al. 2014). Alarmingly, there are only few agents in clinical development for the treatment of infections caused by multidrug-resistant Gram-negative pathogens (Bush and Page 2017).

*Pseudomonas aeruginosa*, the causative agent of severe acute as well as of chronic persistent infections, is particularly problematic. The opportunistic pathogen exhibits high intrinsic antibiotic resistance and frequently acquires resistance conferring genes via horizontal gene transfer (Lister, Wolter, and Hanson 2009; Partridge et al. 2018). Furthermore, the accelerating development of drug-resistance due to the acquisition of drug resistance-associated mutations poses a serious threat.

The lack of new antibiotic options underscores the need for optimization of current diagnostics. Diagnostic tests are a core component in modern healthcare practice. Especially in the light of rising multidrug-resistance, high-quality diagnostics becomes increasingly important. However, to provide information as the basis for infectious disease management is a difficult task. Antimicrobial susceptibility testing (AST) has experienced little change over the years. It still relies on culture-dependent methods and as a consequence, clinical microbiology diagnostics is labor-intensive and slow. Culture-based AST requires 48h (or longer) for definitive results, which leaves physicians with uncertainty about the best drugs to prescribe to individual patients. This delay also contributes to the spread of drug-resistance (López-Causapé et al. 2018; Oliver et al. 2015).

The introduction of molecular diagnostics could become an alternative to culture-based methods and could be critical in paving the way to fight antimicrobial resistance. Identification of genetic elements of antimicrobial resistance promises a deeper understanding of the epidemiology and mechanisms of resistance and could lead to a timelier reporting of the resistance profiles as compared to conventional culture-based testing. It has been demonstrated that for a number of bacterial species antimicrobial resistance can be highly accurately predicted based on information derived from the genome sequence (Gordon et al. 2014; Bradley et al. 2015; Moradigaravand et al. 2018). However, in the opportunistic pathogen *P. aeruginosa* even full genomic sequence information is insufficient to predict antimicrobial resistance in all clinical isolates (Kos et al. 2015). *P. aeruginosa* exhibits a profound phenotypic plasticity mediated by environment-driven flexible changes in the transcriptional profile (Dötsch et al. 2015). For example, *P. aeruginosa* adapts to the presence of antibiotics with the over-expression of the *mex* genes, encoding the antibiotic extrusion machineries MexAB-OprM, MexCD-OprJ, MexEF-OprN and MexXY-OprM. Similarly, high expression of the *ampC* encoded intrinsic beta-lactamase confers antimicrobial resistance (Martin et al. 2018; Goli et al. 2018; Haenni et al. 2017; Juan et al. 2017). Those transcriptional responses are frequently fixed in clinical *P. aeruginosa* strains, e.g. due to mutations in negative regulators of gene expression (Frimodt-Møller et al. 2018; Juarez et al. 2018). Thus, the isolates develop an environment-independent resistance phenotype. Up-regulation of intrinsic beta-lactamases as well as over-expression of efflux pumps that contribute to the resistance phenotype make gene-based testing a challenge, because it is difficult to predict from the genomic sequence, which (combinations of) mutations would lead to an upregulation of resistance conferring genes (Llanes et al. 2004; Schniederjans, Koska, and Häussler 2017; Fernández and Hancock 2012).

In this study, we investigated whether we can reliably predict antimicrobial resistance in *P. aeruginosa* using not only genomic but also quantitative gene expression information. For this purpose, we sequenced the genomes of 414 drug-resistant clinical *P. aeruginosa* isolates and recorded their transcriptional profiles. We built predictive models of antimicrobial susceptibility/resistance to four commonly administered antibiotics by training machine learning classifiers. From these classifiers we inferred candidate marker panels for a diagnostic assay by selecting resistance- and susceptibility-informative markers via feature selection. We found that the combined use of information on the presence/absence of genes, their sequence variation and gene expression profiles can predict resistance and susceptibility in clinical *P. aeruginosa* isolates with high or very high sensitivity and predictive value.

## Results

### Taxonomy and antimicrobial resistance distribution of 414 DNA and mRNA sequenced clinical *P. aeruginosa* isolates

414 *P. aeruginosa* isolates were collected from clinical microbiology laboratories of hospitals across Germany and at sites in Spain, Hungary and Romania (Figure 1A). For all isolates, the genomic DNA was sequenced and transcriptional profiles were recorded. This enabled us to not only use the full genomic information but also information on the gene expression profiles as an input to machine learning approaches.

**Figure 1:**
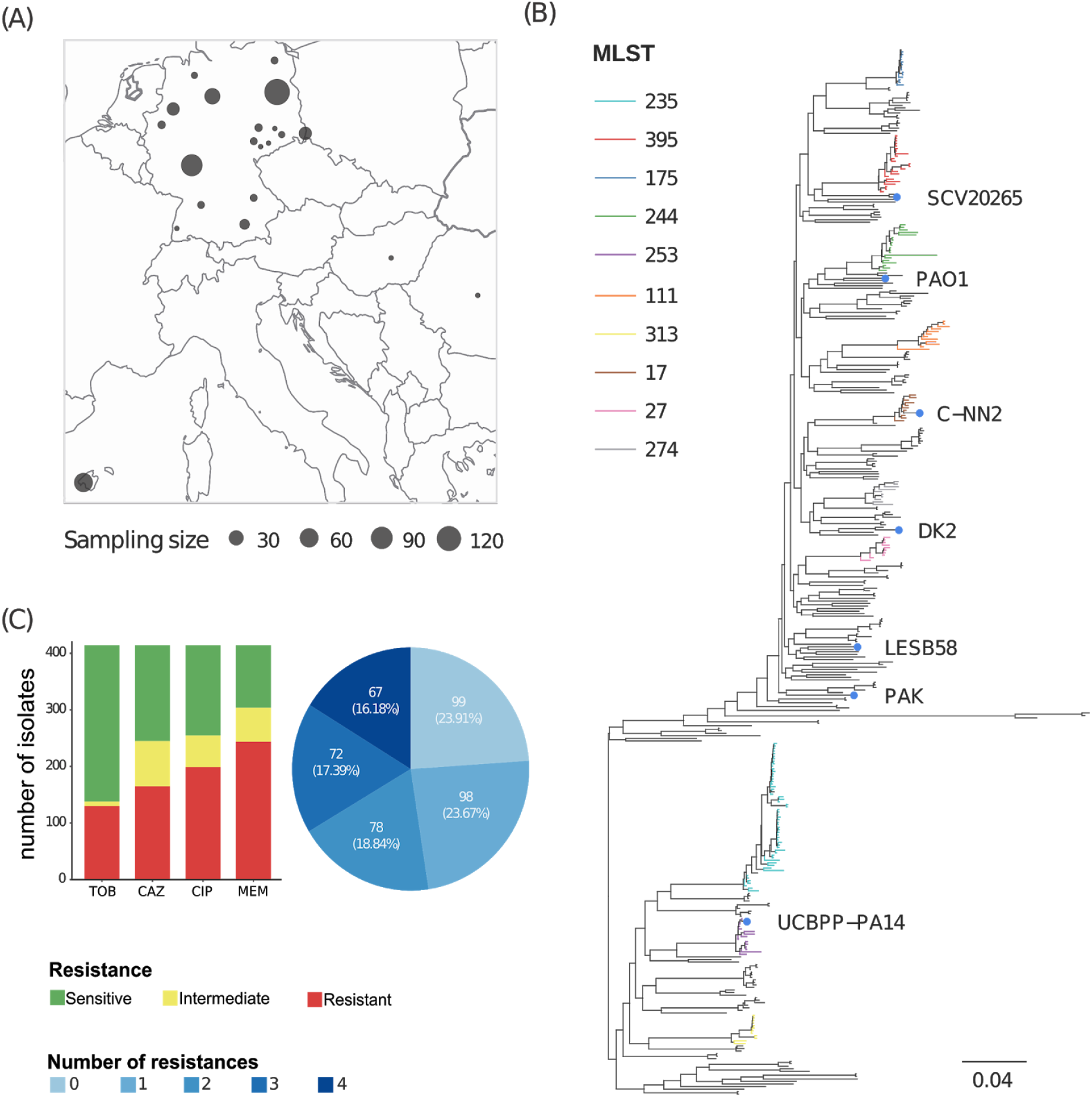
Geographic and phylogenetic distribution of 414 clinical *P. aeruginosa* isolates used in this study. (A) Geographic sampling site distribution, where circle size is proportional to the number of isolates from a particular location. (B) Phylogenetic tree of the clinical isolates and seven reference strains (a PA7-like outgroup clade including two clinical isolates is not shown). Abundant high risk clones identified by multi-locus sequence typing (MLST) are indicated by branch coloring. (C) Antimicrobial susceptibility profile regarding the four commonly administered antibiotics tobramycin (TOB), ceftazidime (CAZ), ciprofloxacin (CIP) and meropenem (MEM) tested by Agar dilution according to Clinical & Laboratory Standards Institute guidelines (CLSI 2018).

We inferred a maximum likelihood phylogenetic tree based on variant nucleotide sites (Figure 1B). The tree was constructed by mapping the sequencing reads of each isolate to the genome of the *P. aeruginosa* PA14 reference strain and then aligning the consensus sequences for each gene. The isolates exhibited a broad taxonomic distribution and separated into two major phylogenetic groups. One included PAO1, PACS2, LESB58, and a cluster of high-risk clone ST175 isolates; the other included PA14, as well as one large cluster of high-risk clone ST235 isolates. Both groups comprised several further clades with closely related isolates of the same sequence type as determined by multi-locus sequencing typing (MLST).

Next, we recorded antibiotic resistance profiles for all isolates regarding the four common anti-pseudomonas antimicrobials, tobramycin (TOB), ceftazidime (CAZ), ciprofloxacin (CIP), and meropenem (MEM) using agar dilution method. Most isolates of our clinical isolate collection exhibit antibiotic resistance against these four antibiotics (Figure 1C, Supplementary Table S1). One third had a multidrug resistant (MDR) phenotype, defined as non-susceptible to at least three different classes of antibiotics (Magiorakos et al. 2012).

### Machine learning for predicting antimicrobial resistance

We used the genomic and transcriptomic data of the clinical *P. aeruginosa* isolates to infer resistance and susceptibility phenotypes to ceftazidime, meropenem, ciprofloxacin and tobramycin with machine learning classifiers. For each antibiotic we included all respective isolates categorized as either “resistant” or “susceptible”. For the genomic data, we included sequence variations (short nucleotide polymorphisms; SNPs, including small indels), and gene presence or absence (GPA) as features. In total, we analysed 255,868 SNPs, represented by 65,817 groups with identical distributions of SNPs across isolates for the same group, and 76,493 gene families with presence or absence information, corresponding to 14,700 groups of identically distributed gene families. 1,306 of these gene families had an indel in some isolate genomes, which we included as an additional feature. We evaluated SNP and GPA groups in combination with gene expression information for 6,026 genes (Figure 2). For each drug we randomly assigned isolates to a training set that comprised 80% of the resistant and susceptible isolates, respectively, and the remaining 20% to a validation set. Parameters of machine learning models were optimized on the training set and their value assessed in cross-validation (CV), while the validation set was used to obtain another independent performance estimate. We trained several machine learning classification methods on SNPs, GPA and expression features individually and in combination for predicting antibiotic susceptibility or resistance of isolates and evaluated the classifier performances. As the underlying gold standard for evaluation, we determined MIC (minimal inhibitory concentration) values of all clinical isolates with agar dilution according to CLSI guidelines (CLSI 2018).

**Figure 2:**
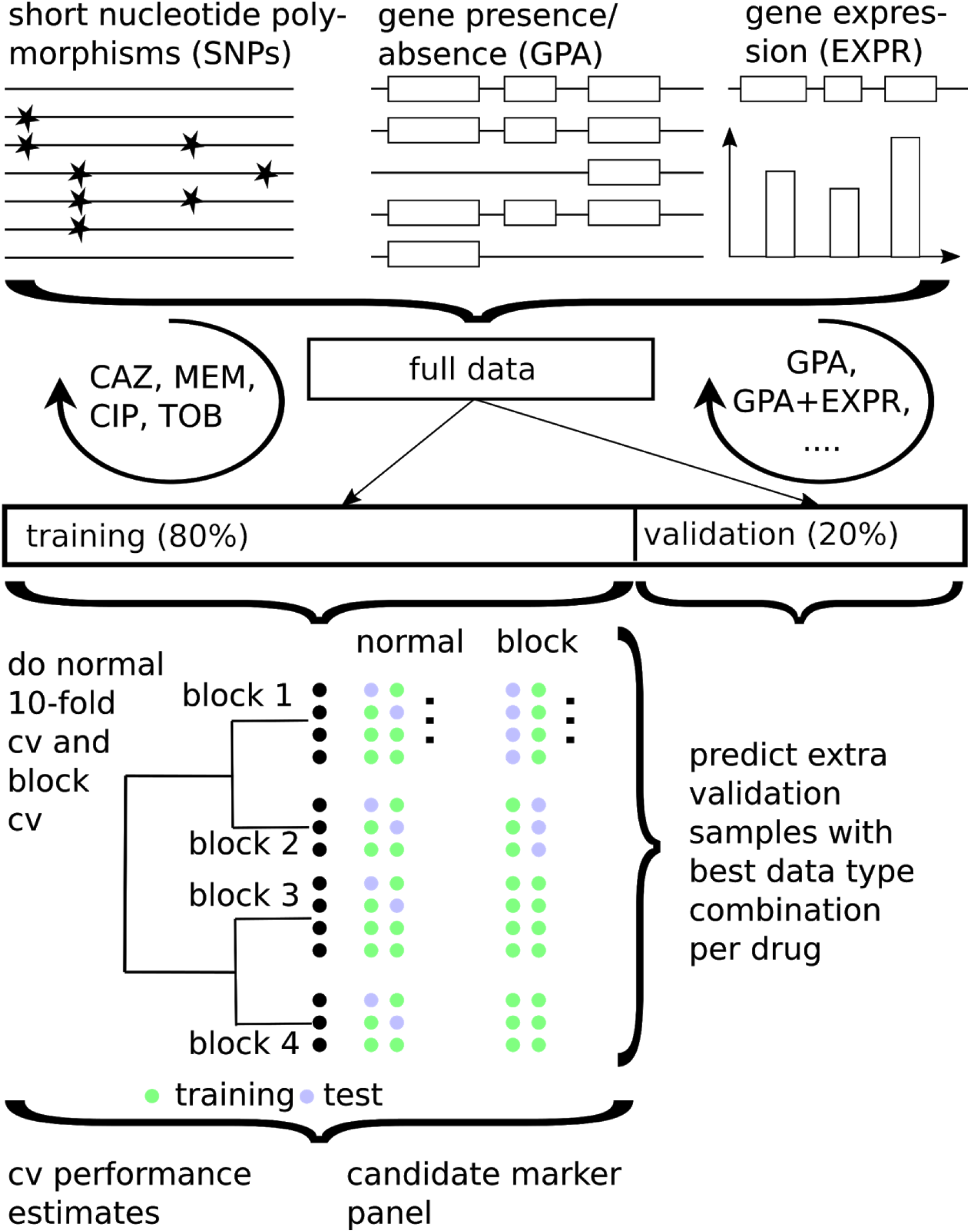
Training and validating a diagnostic classifier for antimicrobial susceptibility prediction for four different drugs based on genomic (GPA/SNPs) and transcriptomic profiles (EXPR). The best data type combination was determined using 80% of the data in standard and phylogenetically-informed cross-validation (cv) and further validated on the remaining 20% of the data.

We calculated the sensitivity and predictive value of resistance (R) and susceptibility (S) assignment, as well as the macro F1-score, as an overall performance measure based of a classifier trained on a specific data type combination. The sensitivity reflects how good that classifier is in recovering the assignments of the underlying gold standard, representing the fraction of susceptible, or resistant, samples, respectively. The predictive value reflects how trustworthy the assignments of this particular classifier are, representing the fraction of correct assignments of all susceptible, or resistant assignments, respectively. The F1-score is the harmonic mean of the sensitivity and predictive value for a particular class, i.e. susceptible or resistant. The macro F1-score is the average over the two F1-scores.

We used the support vector machine (SVM) classifier with a linear kernel, as in (Weimann et al. 2016), to predict sensitivity or resistance to four different antibiotics. Parameters were optimized in nested CV and performance estimates averaged over five repeats of this setup. The combined use of i) GPA, ii) SNPs and iii) information on gene expression resulted in high (0.8-0.9) or very high (>0.9) sensitivity and predictive values (Figure 3). Notably, the relative contribution of the different information sources to the susceptibility and resistance sensitivity strongly depended on the antibiotic. To assess the effect of the classification technique, we compared the performance of an SVM classifier with a linear kernel to that of random forests, and logistic regression, which we and others have used successfully for related phenotype prediction problems (Her and Wu 2018; Asgari et al. 2018; Wheeler, Gardner, and Barquist 2018). For this purpose we used the data type combination with the best macro F1-score in resistance prediction with the SVM. As before, we evaluated the classification performance in random cross-validation, and on a held-out test data set. In addition, we performed a phylogeny-aware partitioning of our data set, to assess phylogenetic generalization ability of our technique.

**Figure 3:**
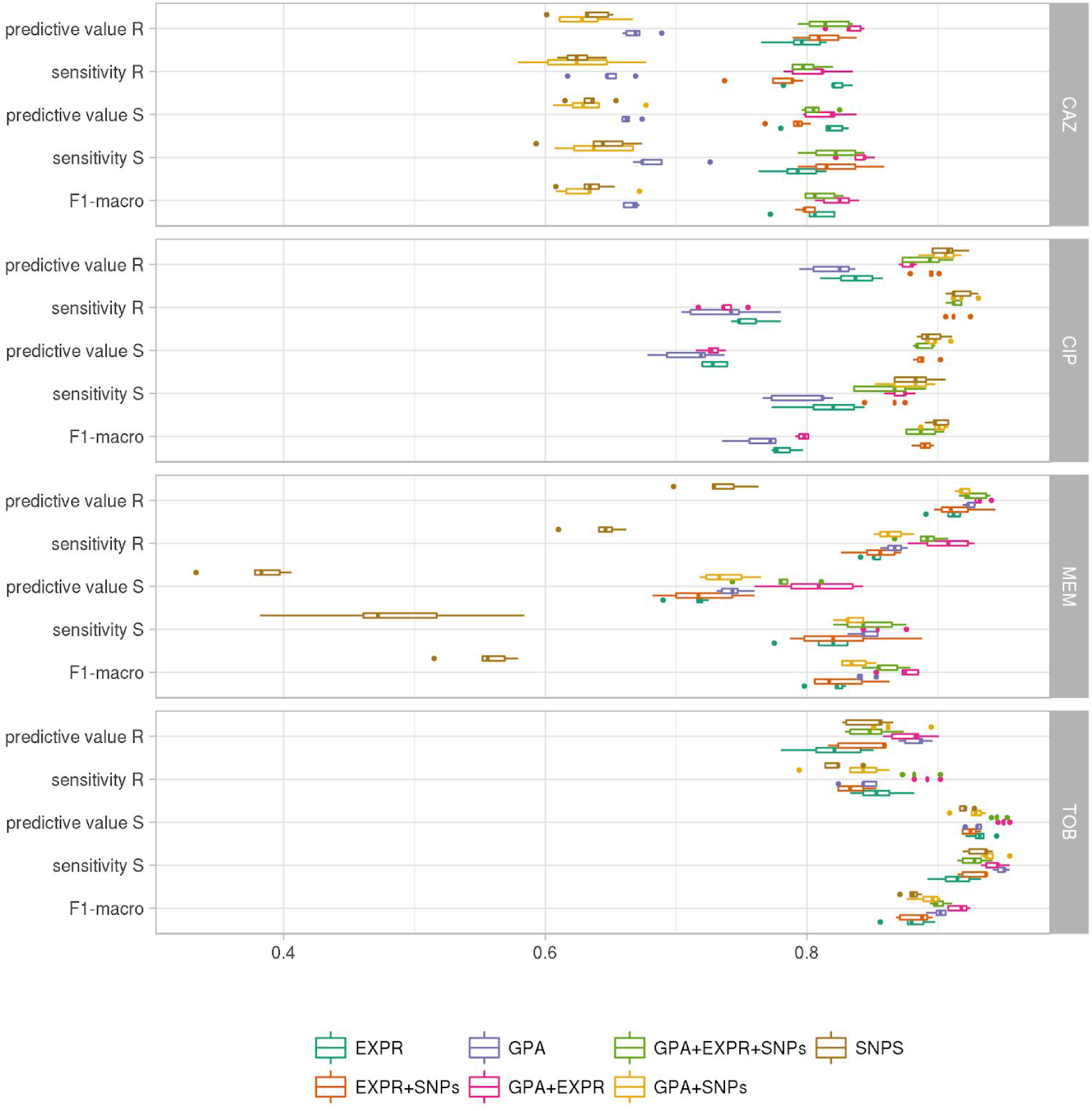
Evaluation of AMR classification with a support vector machine (R: resistant; S: susceptible) using different performance metrics and data types (EXPR: gene expression; GPA: gene presence or absence; SNPs: single nucleotide polymorphisms) or combinations thereof. Each individual panel depicts the results for one of four different anti-pseudomonal antibiotics (CAZ, CIP, MEM, TOB).

The performance of the SVM in random cross-validation was comparable to logistic regression (Macro F1-score for the SVM: 0.83±0.06 vs. logistic regression: 0.84±0.06), but considerably better than the random forest classifiers (0.67±0.14) (Supplementary Figures S1, S2, Supplementary Table S2). The performance on the held-out data set was in a comparable range (SVM: 0.87±0.07, logistic regression: 0.90±0.04, random forest 0.71 ±0.16). We furthermore observed similar macro F1-scores inferred in the phylogenetically selected cross-validation (SVM: 0.87±0.07, logistic regression: 0.86±0.07, random forest 0.72±0.13), which suggests only a minor influence of the bacterial phylogeny on the classification performance. The performance on the phylogenetically selected held-out data set were again comparable, though performance for the random forest deteriorated in comparison to the cross-validation results (SVM: 0.86±0.06, logistic regression 0.83±0.06, random forests 0.56±0.03).

Ciprofloxacin resistance and susceptibility based on SVMs could be correctly predicted with a sensitivity of 0.92±0.01 and 0.87±0.01, and with simultaneously high predictive values of 0.91 ±0.01 and 0.90±0.01, respectively, using solely SNP information. The sensitivity 0.80±0.04 and 0.79±0.02 and predictive value 0.73±0.01 and 0.76±0.02 to predict ciprofloxacin susceptibility and resistance based exclusively on gene expression data was also high. However, there was no added value of using information on gene expression in addition to SNP information for the prediction of susceptibility/resistance towards ciprofloxacin.

For the prediction of tobramycin susceptibility and resistance, the machine learning classifiers performed almost equally well when the three input data types (SNPs, GPA and gene expression) were used individually (values >0.8). SNP information was predictive of tobramycin resistance; however, it did not further improve the classification performance when combined with the other data types. GPA information alone was the most important data type for classifying tobramycin resistance and susceptibility providing sensitivity values of 0.84±0.01 and 0.95±0.01 and predictive values of 0.88±0.01 and 0.93±0.01, respectively. The performance of GPA-based prediction increased further when gene expression values were included (sensitivity values of 0.89±0.01 and 0.94±0.01 for resistance and susceptibility prediction, respectively, and predictive values of 0.88±0.01 and 0.95±0.01).

For the correct prediction of meropenem resistance/susceptibility, gene presence/absence was most influential (sensitivity values of 0.87±0.01 and 0.84±0.01 for resistance and susceptibility prediction, respectively, and predictive values of 0.92±0.00 and 0.74±0.01). As observed for tobramycin, the use of genome-wide information on GPA, and of information on gene expression in combination increased the sensitivity to detect resistance as well as susceptibility to meropenem to 0.91 ±0.02 and 0.86±0.01 and the predictive values to 0.93±0.01 and 0.81 ±0.03, respectively.

For ceftazidime using only information on gene presence/absence revealed a sensitivity of susceptibility/resistance prediction of 0.69±0.01 and 0.66 ±0.01, and predictive values of 0.66±0.01 and 0.67±0.01, respectively. Adding gene expression information considerably improved the performance of susceptibility and resistance sensitivity to 0.83±0.02 and 0.81 ±0.02 and predictive values of 0.81 ±0.02 and 0.83±0.01. In summary, for tobramycin, ceftazidime and meropenem combining GPA and expression information gave the most reliable classification results, whereas for ciprofloxacin we found that only using SNPs provided the best performance (Table 1 and Supplementary Table S3). Thus for the remainder of the manuscript, we will focus on the results obtained with classifiers trained on those data type combinations.

**Table 1:** Performance of support vector machine (SVM) classifier to predict sensitivity or resistance to four different antibiotics. The last column indicates the number of (combined) features that resulted in the least complex SVM model within one standard deviation of the peak performance, i.e. with the best F1-score macro and as few as possible features for each drug.

| Anti-biotic | Markers used | Sensitivity resistance | Sensitivity susceptibility | Predictive value resistance | Predictive value susceptibility | F1- score | Number of markers |
| --- | --- | --- | --- | --- | --- | --- | --- |
| CAZ | GPA+ EXPR | 0.83±0.02 | 0.81±0.02 | 0.81±0.02 | 0.83±0.01 | 0.82±0.01 | 37 |
| TOB | GPA+ EXPR | 0.89±0.01 | 0.94±0.01 | 0.88±0.01 | 0.95±0.01 | 0.92±0.01 | 59 |
| MEM | GPA+ EXPR | 0.91±0.02 | 0.86±0.01 | 0.93±0.01 | 0.81±0.03 | 0.87±0.01 | 93 |
| CIP | SNPs | 0.92 ± 0.01 | 0.87 ± 0.01 | 0.91±0.01 | 0.90±0.01 | 0.90±0.01 | 50 |

### A candidate drug resistance marker panel

We determined the minimal number of molecular features required to obtain the highest macro F1-score for each drug. We inferred the number of features contributing to the classification from the number of non-zero components of the SVM weight vectors, using a standard cross-validation set-up. For each value of the C parameter, which controls the amount of regularization imposed on the model, the cross-validation procedure was repeated five times (Figure 4, Supplementary Table S4). Performance of antimicrobial resistance prediction peaked for the candidate classifiers using between 50 and 100 features. Notably, the ciprofloxacin classifier required only two SNPs until the learning curve performance was almost saturated, whereas classifiers of drugs that included expression and gene presence/absence markers required more feature (>50) to reach saturation.

**Figure 4:**
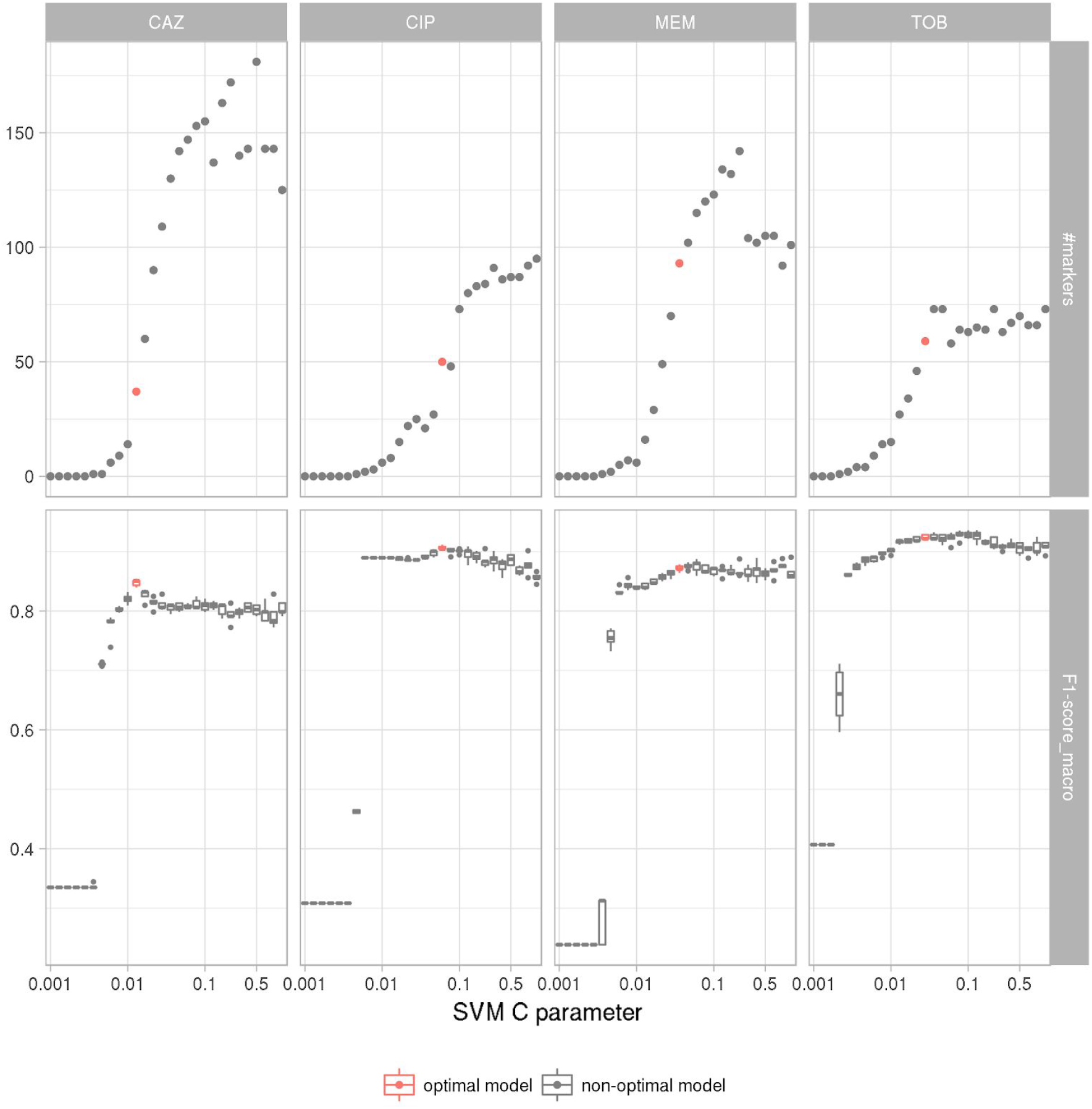
The number of features used by the support vector machine classifier (top panels) and corresponding classification performance (bottom panels) varies with the hyperparameter C. The SVM resistance/susceptibility classifier was evaluated in five repeats of ten-fold nested cross-validation. Each panel depicts the results for a different drug (CAZ, CIP, MER, TOB) based on the best data type combination (GPA+EXPR/SNPs). The model with the fewest features within one standard deviation of the maximal performance was selected as the most suitable diagnostic classification model (red) (Supplementary Table S5).

Next, we determined the C parameter resulting in the least complex SVM model within one standard deviation of the peak performance, i.e. with the best F1-score macro and as few as possible features for each drug (Friedman, Hastie, and Tibshirani 2001). We chose our candidate marker panel for each drug as the set of all non-zero features, and designated the respective model as the most suitable diagnostic classifier. We used SNP information for ciprofloxacin resistance and susceptibility prediction and the combination of GPA and expression features for tobramycin, meropenem and ceftazidime. We refer to each of these classifiers as the candidate classifier for susceptibility and resistance prediction for a particular drug. The ciprofloxacin candidate marker panel contained 50 SNPs. The meropenem, ceftazidime and tobramycin marker lists consisted of 93, 37 and 59 expression and GPA features. The complete list of candidate markers for the prediction of resistance against the four antibiotics is given in Supplementary Table S5. This list includes the candidate markers of the three input features namely GPA, gene expression and SNPs alone and in combination. Table 2 is a shortlist of the panel markers for each drug based on the data combination that had allowed us to train the most reliable classifier.

**Table 2:**
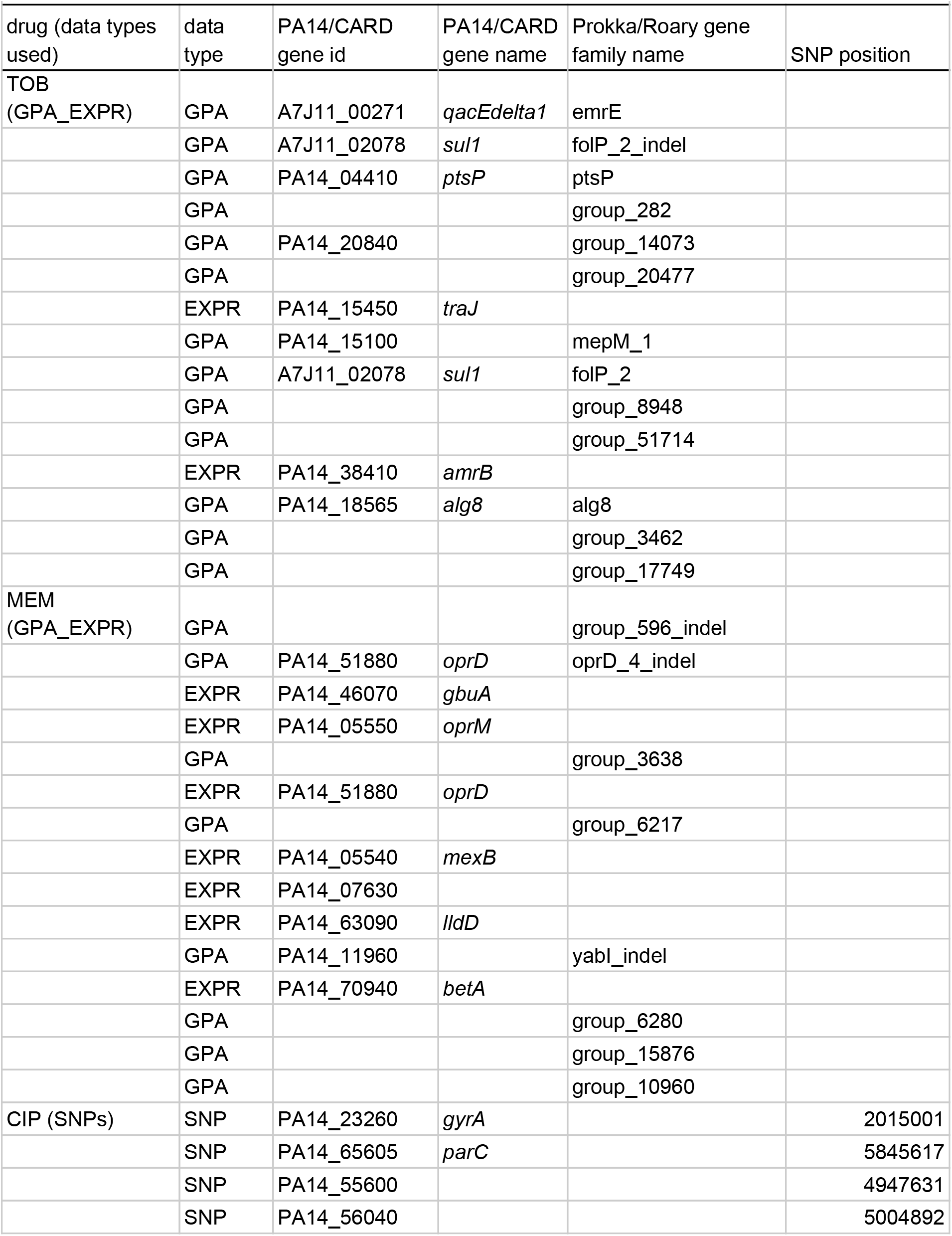
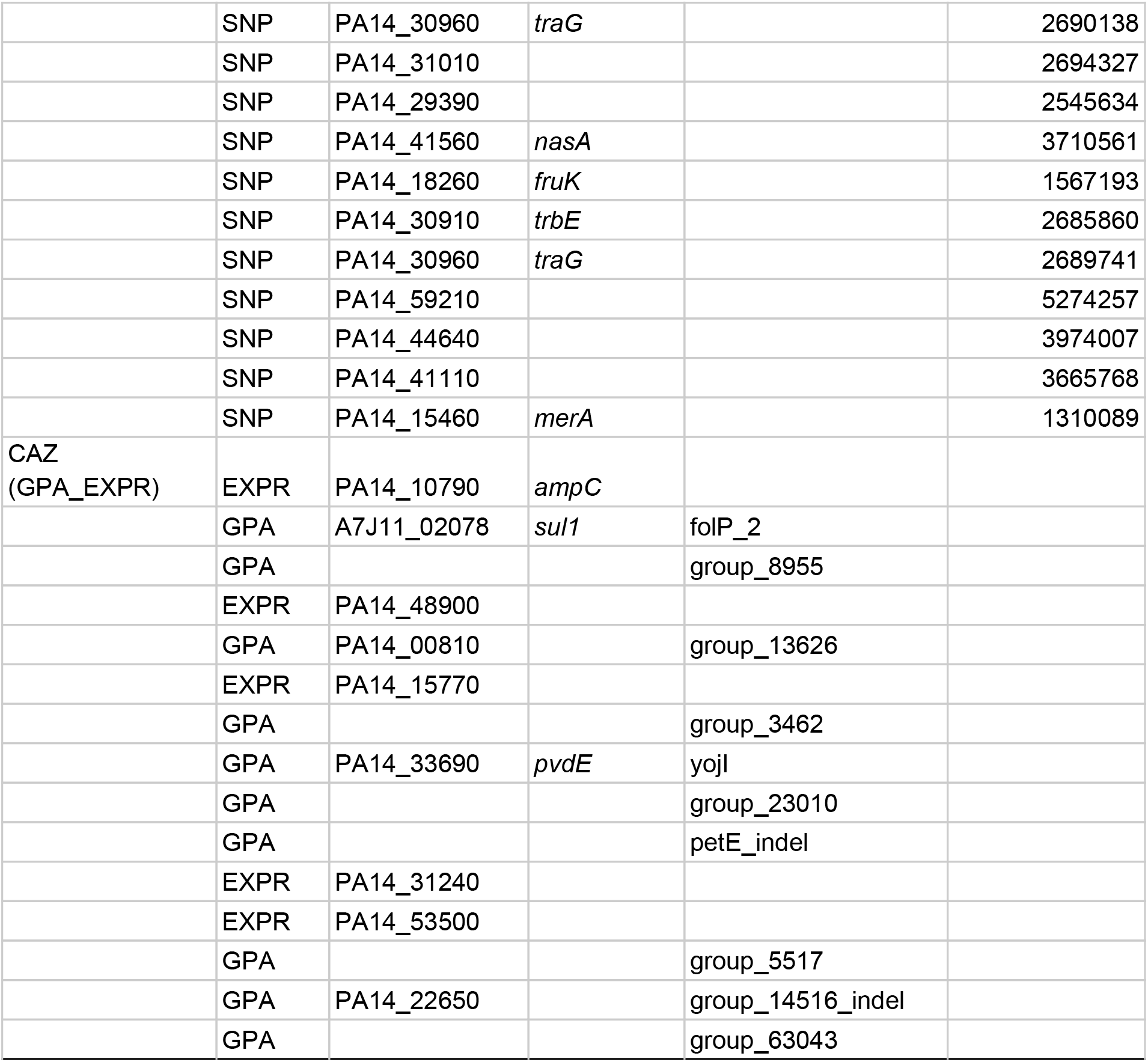
The top 15 candidate markers ranked according to the contribution of each marker to the support vector machine classifier for each drug based on the best performing combination of data types. For gene presence/absence (GPA) markers we provide the gene id and accession based *PA14* reference genome gene family member or based on the Comprehensive Antibiotic Resistance Database (CARD) (Jia et al. 2017). Otherwise we include the gene name or id of each marker as generated by the bacterial genome annotation tool Prokka (Seemann 2014) and protein family clustering software Roary (Page et al. 2015). Expression markers are based on the *PA14* genome, too. For short nucleotide polymorphisms (SNPs) we report the genome position in the reference *PA14* genome.

To test the performance of the candidate marker panel-based classifiers on an independent set of clinical *P. aeruginosa* isolates, we used them to predict antibiotic resistance for the samples of the validation dataset (Figure 5, Supplementary Table S6). On this held out data we obtained an F1-sore for all drugs that was similarly high as before: namely this was for meropenem 0.95, ceftazidime 0.77 and tobramycin 0.96, using gene expression and gene presence/absence features and 0.87 for ciprofloxacin using SNP information. These results indicate that the diagnostic classifiers have good generalization abilities when applied to new samples. We observed more variability across drugs than in nested cross-validation, which is expected due to the smaller size of the validation set.

**Figure 5:**
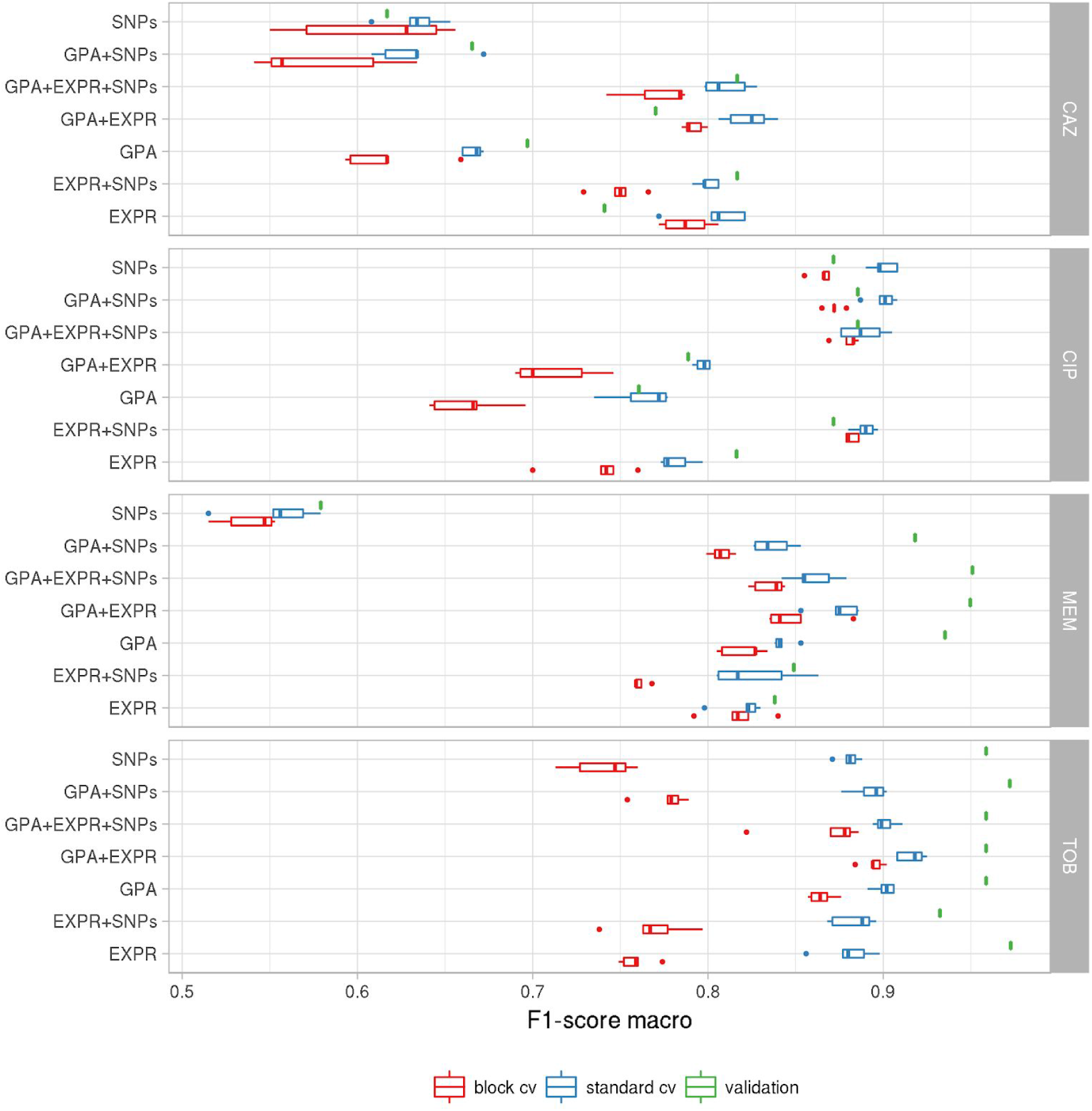
Performance of the support vector machine (SVM) classifier for antimicrobial resistance and susceptibility prediction for different data types, different drugs and different evaluation schemes. The SVM performance was summarized by the F1-score and is shown for standard cross-validation (standard_cv, blue) and cross-validation using phylogenetically related blocks of isolates (block_cv, red) based on the training data set (80% of the isolates) and for the validation data set (green; 20% of the isolates). EXPR: gene expression; GPA: gene presence and absence with indel information. SNPs: short nucleotide polymorphisms.

### Improvement of assignment accuracy with increasing sample numbers

We next investigated how prediction performance depended on the number of samples used for classifier training. We trained the SVM classifiers on random subsamples of different sizes of the full data set with 414 isolates. For each model we recorded the macro F1-score in five repeats of ten-fold nested cross-validation (Figure 6). The classification performance saturates for all our classifiers well before using all available training samples, suggesting that when adding more isolates for resistance classification, the classification performance would improve only very slowly. Markers potentially remaining undiscovered in our study might have very small effect sizes, requiring much larger data set sizes for their detection. Interestingly, the number of samples required until the performance curve plateaued depends on the drugs and data types used. For ciprofloxacin, the performance of susceptibility/resistance prediction based on SNPs saturated quickly, likely due to the large impact of the known mutations in the quinolone resistance determining region (QRDR), whereas the classifiers for the other three drugs, which were trained on expression and gene presence/absence information required more samples until the F1-score plateaued. For these classifiers the dispersion of the macro F1-score for subsets of the data with fewer samples is also considerably higher than for the ciprofloxacin SNP models.

**Figure 6:**
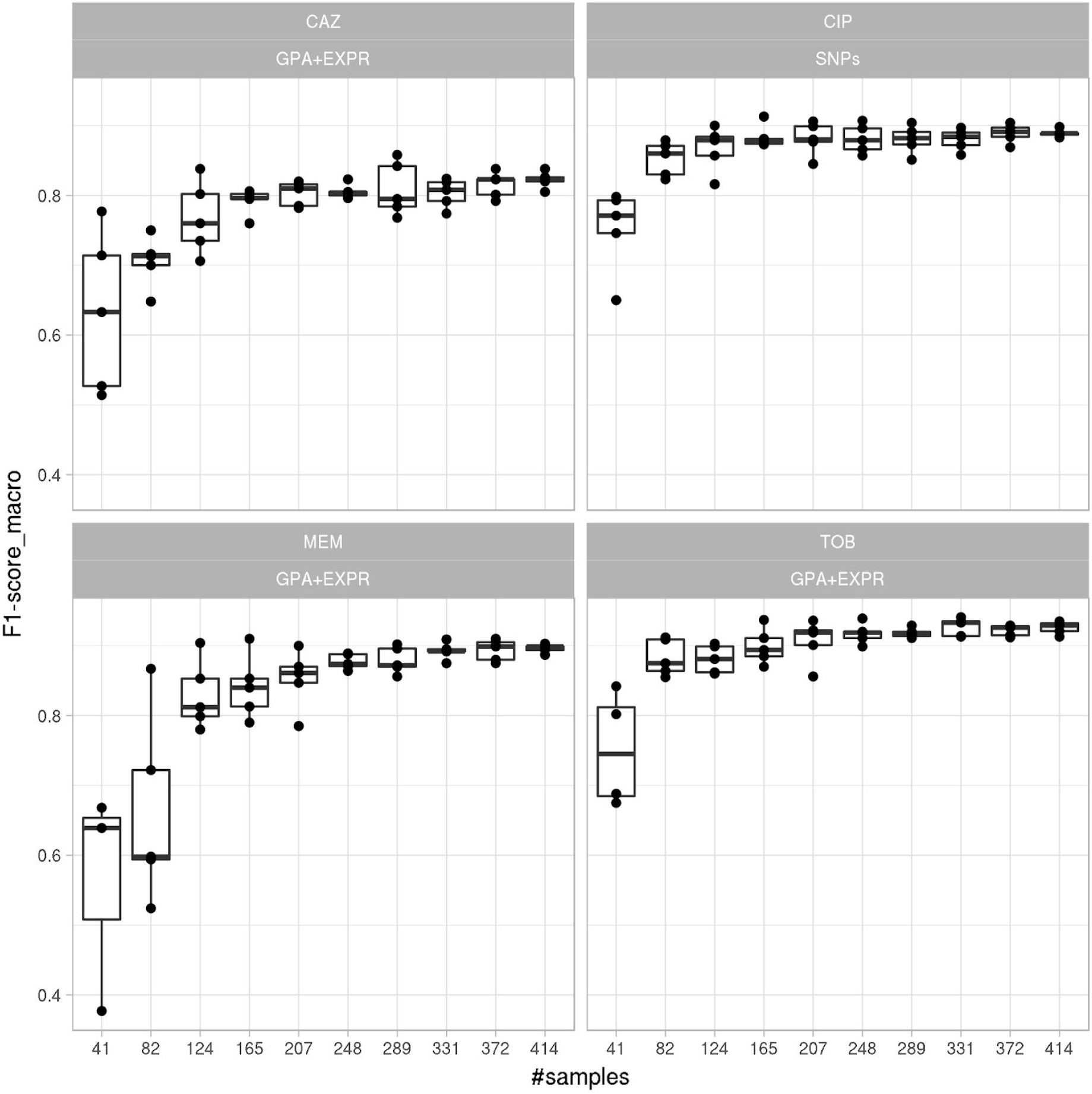
Classification performance improves and plateaus with the number of training samples used. A support vector machine-based resistance/susceptibility classifier was trained on differently sized and randomly drawn sub-samples from our isolate collection and evaluated in five repeats of a ten-fold nested cross-validation. Each panel depicts the results for a different drug (CAZ, CIP, MEM, TOB) based on the best data type combination (GPA+EXPR/SNPs).

### Performance estimates of the candidate diagnostic classifiers are independent of the bacterial phylogeny

In *P. aeruginosa*, different phylo-groups might contain different antibiotic resistance genes or mutations alone or in combinations. Thus, if there was an association of distinct resistance conferring genes with certain phylo-groups, our machine learning approach might identify markers that distinguish between different phylo-groups rather than between susceptible and resistant clinical isolates. In Supplementary Figures 3, 4, 5 and 6 we show susceptibility and resistance of each isolate in the context of the phylogenetic tree as predicted by the diagnostic classifier and based on AST for each of the drug. To assess whether our predictive markers are biased by the phylogenetic structure of the clinical isolate collection, we assessed classification robustness in a block cross-validation approach. Here, isolates of phylo-groups with differing sequence types as determined by MLST were grouped into blocks and all isolates of a given block were only allowed to be either in the training or test folds (Figure 2, 5). Overall, for all four candidate diagnostic classifiers, we found that the performance estimates were only slightly lower than those obtained with standard cross-validation. This confirmed that the various *P. aeruginosa* phylogenetic subgroups possess similar mechanisms and molecular markers for the resistance phenotype and that the identified markers are distinctive for resistance/susceptibility instead of phylogenetic relationships.

Notably, this was different for some suboptimal data type combinations, such as for predicting tobramycin resistance using SNPs or gene expression, where a substantially lower discriminative performance was achieved in block-compared to random cross-validation (macro F1-score difference > 0.2, Supplementary Table S3). These results suggest that – despite the observed independence of the presence of genetic resistance markers and bacterial phylogeny – for some antibiotics we found a non-negligible phylo-group-dependent performance effect. This underlines the importance of assessing the impact of the phylogeny on the antimicrobial resistance prediction.

### Misclassified isolates are more frequent near the MIC breakpoints

We tested whether we could detect an overrepresentation of misclassified samples among the samples with a MIC value close to the breakpoints compared to samples with higher or lower MIC values, selecting samples from equidistant intervals (in log space) around the breakpoint. We report only the strongest overrepresentation for each drug after multiple testing correction. For ciprofloxacin significantly more samples with a MIC between 0.5 and 8 were misclassified (31 of 139 samples (22%)) than samples with a MIC smaller than 0.5 or larger than 8 (7 of 219 samples (3%)) (Fisher exact test with an FDA adjusted p-value of 6.2·10^-8^; Figure 7). For ceftazidime, we found that 46 of 177 samples (26%) with a MIC between 4 and 64 were misclassified whereas only 21 of 157 (13%) of samples with a MIC smaller or higher than those values were misclassified (adjusted p-value: 0.014). For meropenem, we found that 26 of 207 samples (13%) with a MIC between 1 and 16 were misclassified, but only 8 of 147 (5%) of all samples with a MIC smaller or higher than those values were misclassified (adjusted p-value: 0.05). For tobramycin, no significant difference was found.

**Figure 7:**
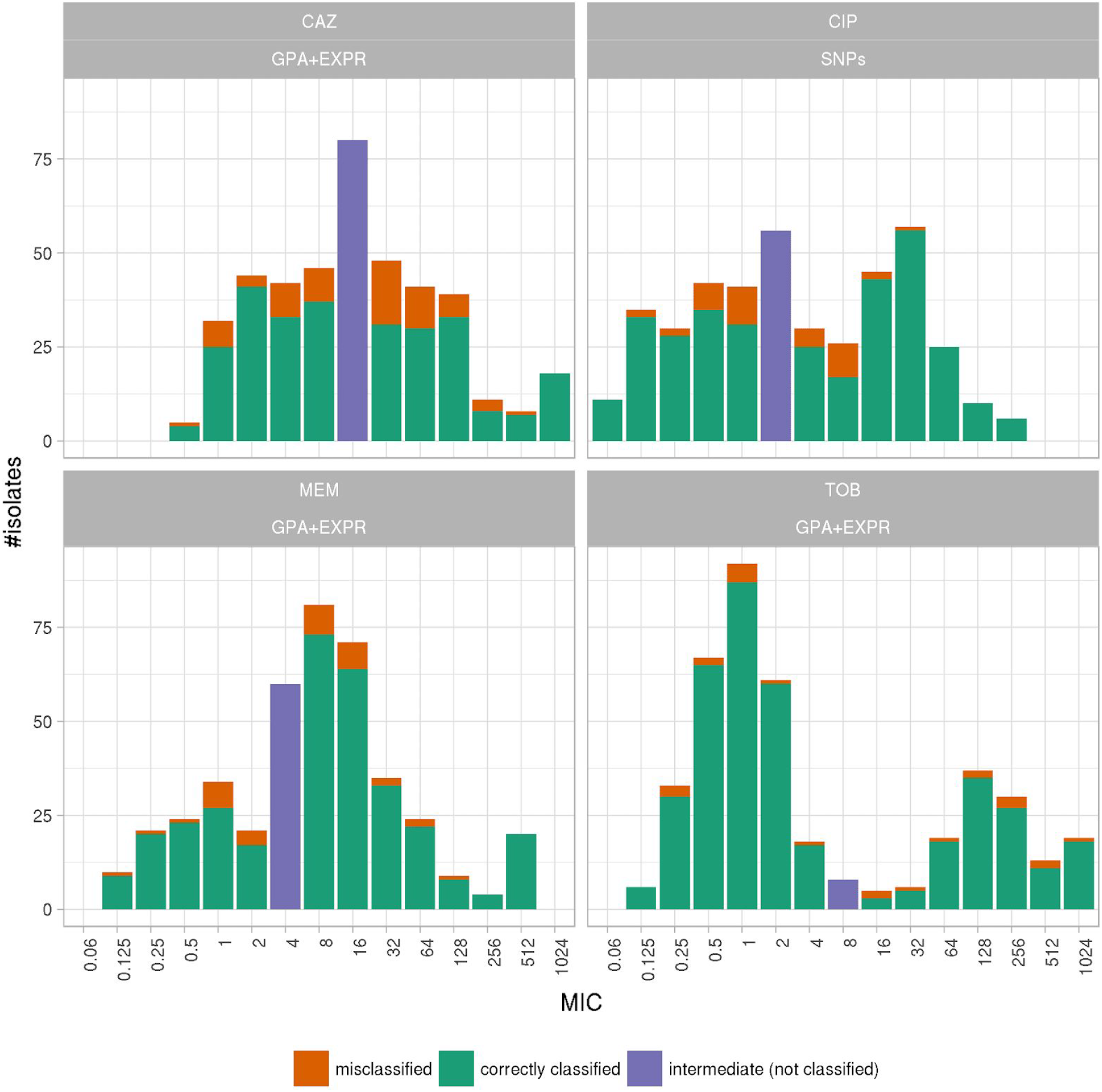
Number of samples misclassified and correctly predicted by the support vector machine resistance and susceptibility classifier (SVM) grouped by their minimum inhibitory concentration. Each panel depicts the results for a different anti-pseudomonal drug (CAZ: ceftazidime, CIP: ciprofloxacin, MEM: meropenem, TOB: tobramycin) for the best data type combination (GPA+EXPR/SNPS). Misclassified and correctly classified samples for the training data set (80%) were inferred in a ten-fold cross-validation. An SVM trained on the training data set was used to predict resistance/susceptibility of the validation samples (20%). The number of misclassified samples in the training (80%) and validation set were aggregated.

## Discussion

One of the most powerful weapons in the battlefield of drug-resistant infections is rapid diagnostics of resistance. Earlier and more detailed information on the pathogens antimicrobial resistance profile has the potential to change antimicrobial prescribing behavior and improve the patient’s outcome. The demand for faster results has initiated investigation of molecular alternatives to today’s culture-based clinical microbiology procedures. However, for the successful implementation of robust and reliable molecular tools, it is critical to identify the entirety of the molecular determinants of resistance. Failure to detect resistance can lead to the administration of ineffective or sub-optimal antimicrobial treatment. This has direct consequences for the patient; and poses significant risks especially in the critically ill patient. Conversely, failing to identify susceptibility may result in the avoidance of a drug despite the fact that it would be suitable to treat the pathogen, in the extreme case leading to patient death due to a lack of known treatment options. Overtreatment could also be a consequence and the needless use of broad-spectrum antibiotics. This drives costs in the hospital, puts patients at risk for more severe side effects and may contribute to the development of drug resistance by applying undesired selective pressures.

In this study, we show that without any prior knowledge on the molecular mechanisms of resistance, machine learning approaches using genomic and transcriptomic features can provide high antibiotic resistance assignment capabilities for the opportunistic pathogen *P. aeruginosa*. The performance of drug resistance prediction was strongly dependent on the antibiotic.

Ciprofloxacin resistance and susceptibility prediction mostly relied on SNP information. Particularly two SNPs in the quinolone resistance determining region (QRDR) of *gyrA* and *parC* had the strongest impact on the classification (Supplementary Table S3). This is an expected finding as quinolone antibiotics act by binding to their targets, gyrase and topoisomerase IV (Bruchmann et al. 2013); and target-mediated resistance caused by specific mutations in the encoding genes is the most common and clinically significant form of resistance (del Barrio-Tofiño et al. 2017). Although the sensitivity to predict resistance and susceptibility from only gene expression data was also high towards ciprofloxacin, there was no added value of using information on gene expression in addition to SNP information. Nevertheless, for the design of a diagnostic test system, it might be of value to include also gene expression information as a fail-safe strategy. Interestingly, among the gene expression classifiers that were associated with ciprofloxacin susceptibility/resistance, we found *prtN*, which is involved in pyocyanin production. Pyocyanin has previously been demonstrated to alter ciprofloxacin susceptibility in *P. aeruginosa* (Grant et al. 2010). Furthermore, we found that altered expression and sequence of genes of the conjugation machinery such as *traG, trbE, trbI* and the closely located transcriptional regulator *merD* influenced classification. This is interesting because, it has been shown before that quinolone resistance inducing QRDR mutations do not only appear as a result of *de novo* target modification but are also frequently obtained by conjugation (Pitondo-Silva et al. 2015).

For the prediction of tobramycin susceptibility and resistance, the machine learning classifiers performed almost equally well when the three input data types (SNPs, GPA and gene expression) were used individually (sensitivity and predictive values >0.8). Remarkably, the combined use of the GPA and the gene expression data sets improved the classification performance. Although SNP information also was predictive of tobramycin resistance, it did not further improve the classification performance when combined with the other feature types. GPA information alone was the most important data type for classifying tobramycin resistance or susceptibility. The majority of aminoglycoside resistant clinical isolates harbor genes encoding for aminoglycoside modifying enzymes (AMEs). The AMEs are very diverse but are usually encoded by genes located on mobile genetic elements, including integrons and transposons. In accordance, the presence of respective markers that indicate the presence of these mobile elements was found to be strongly associated with tobramycin resistance (e.g. *qacE, delta1, sul1* or *folP*). However, the most influential discriminator was the presence of the *emrE* gene. *EmrE* has been described to directly impact on aminoglycoside resistance by mediating the extrusion of small polyaromatic cations (X.-Z. Li, Poole, and Nikaido 2003). Expression of *ptsP* was also associated with tobramycin resistance. This gene has previously already been associated with tobramycin resistance (Schurek et al. 2008). The performance of GPA-based prediction increased further when gene expression values were included. We found e.g. *amrB* (*mexY*), which encodes a multidrug efflux pump known to confer to aminoglycoside resistance (Westbrock-Wadman et al. 1999; Lau, Hughes, and Poole 2014), as one of the top candidates within the marker panel. This confirms that expression of efflux pumps is an important bacterial trait that drives the resistance phenotype in *P. aeruginosa*. Tobramycin resistance/susceptibility was associated especially with an altered expression or SNPs within genes involved in type 4 pili motility (*pilB pilT2, pilY1, pilQ, pilH*) and the type three secretion system (*pcr* genes). It has been proposed that surface motility can lead to extensive multidrug adaptive resistance as a result of the collective dysregulation of diverse genes (Sun et al. 2018). Furthermore, the expression of the type III secretion effector gene, *exoU*, has been previously shown to be associated with antibiotic resistance to tobramycin in *P. aeruginosa* strains isolated from chronic otitis media infections (Park et al. 2017).

For the correct prediction of meropenem resistance/susceptibility, gene presence/absence was most influential. Interestingly, in contrast to tobramycin resistance classification, we observed a substantial accumulation of indels in specific marker genes. Among these marker genes were *ftsY*, involved in targeting and insertion of nascent membrane proteins into the cytoplasmic membrane, *czcD*, encoding a cobalt-zinc-cadmium efflux protein and *oprD*. Inactivation of the porin OprD is the leading cause of carbapenem non-susceptibility in clinical isolates (Köhler et al. 1999). As expected, also a decreased *oprD* gene expression in the resistant group of isolates was identified as an important discriminator. Interestingly though, the most important gene expression marker was not the down-regulated *oprD*, but an up-regulation of the gene *gbuA*, encoding a guanidinobutyrase in the arginine dehydrogenase pathway, in the meropenem resistant group of isolates. The finding of *gbuA* expression in meropenem resistant *P. aeruginosa* isolates could be an interesting topic to further follow up. Furthermore, components encoding the MexAB-OprM efflux pump (*mexB, oprM)* were identified as important features associated with resistance. This efflux pump is known to export beta-lactams, including meropenem (Srikumar et al. 1998; X. Z. Li, Nikaido, and Poole 1995; Clermont, Brahimi, and Plesiat 2001).

As observed for tobramycin, the correct prediction of ceftazidime resistance/susceptibility was strongly influenced by both, gene expression values (here *ampC, fpvA, pvdD, algF*) and gene presence/absence (including presence of mobile genetic elements). AmpC is a known intrinsic beta-lactamase, able to hydrolyze cephalosporins (Lister, Wolter, and Hanson 2009). Adding information on the gene expression considerably improved the performance of susceptibility and resistance sensitivity, which was not observed in a similar scale for any other antibiotic.

In conclusion, we demonstrate that extending the genetic features (SNPs and gene presence/absence) with gene expression values is key to improving performance. Thereby relative contribution of the different categories of biomarkers to the susceptibility and resistance sensitivity strongly depended on the antibiotic. This is in stark contrast to the prediction of antibiotic resistance in many Enterobacteriaceae, where knowledge of the presence of resistance-conferring genes, such as beta-lactamases, is usually sufficient to correctly predict the susceptibility profiles. However, analysis of the gene expression marker lists revealed that the resistance phenotype in the opportunistic pathogen *P. aeruginosa* (and possibly also in other non-fermenters) is multifactorial and that alterations in gene expression can alter the resistance phenotype quite substantially.

Intriguingly, we found that the performance of our classifiers improved if the isolates exhibited MIC values that were not close to the breakpoint. This was especially apparent for ciprofloxacin. It has been demonstrated that patients treated with levofloxacin for bloodstream infections caused by Gram-negative organisms for which MICs were elevated, yet still in the susceptible category, had worse outcomes than similar patients infected with organisms for which MICs were lower (Defife et al. 2009). A possible explanation for treatment failure could be the presence of first-step mutations in *gyrA* that lead to MIC values near the breakpoint. If subjected to quinolones, those isolates can rapidly acquire second step mutations in *parC* that would then exhibit a fully resistant phenotype. An additional explanation might also be that generally MICs have a low level of reproducibility (Turnidge and Paterson 2007; Juan et al. 2012; Javed et al. 2018). A non-accurate categorization due to uncertainty in testing near the MIC breakpoint can explain failure in the assignment of drug resistance by the machine learning classifiers.

Capturing the full repertoire of markers that are relevant for predicting antimicrobial resistance in *P. aeruginosa* will require further studies, to expand the predictive power of the established marker lists. The remaining misclassified samples in our study on the basis of these marker lists represent a valuable resource to uncover further spurious resistance mutations.

The broad use of molecular diagnostic tests promises more detailed and timelier information on antimicrobial resistant phenotypes. This would enable the implementation of early and more targeted, and thus more effective antimicrobial therapy for improved patient care. Importantly, a molecular assay system can easily be expanded to test for additional information such as the clonal identity of the bacterial pathogen or the presence of critical virulence traits. Thus, availability of molecular diagnostic test systems can also provide prognostic markers for disease outcome and give valuable information on the clonal spread of pathogens in the hospital setting. However, to realize the full potential of the envisaged molecular diagnostics, clinical studies will be needed to demonstrate that broad application of such test systems will have an impact in clinical decision-making, provide the basis for more efficient antibiotic use, and also decrease the costs of care.

## Materials and Methods

### Strain collection and antibiotic resistance profiling

Our study included 414 clinical *P. aeruginosa* isolates provided by different clinics or research institutions: 350 isolates were collected in Germany (138 at the Charité Berlin (CH), 89 at the University Hospital in Frankfurt (F), 39 at the Hannover Medical School (MHH), and 84 at different other locations). 62 isolates were provided by a Spanish strain collection located at the Son Espases University Hospital in Palma de Mallorca (ESP), and two samples originated from Hungary and Romania, respectively.

All clinical isolates were tested for their susceptibility towards the four common anti-pseudomonas antibiotics tobramycin (TOB), ciprofloxacin (CIP), meropenem (MEM), and ceftazidime (CAZ). Minimal inhibitory concentration (MIC) testing and breakpoint determination was performed in agar dilution according to Clinical & Laboratory Standards Institute (CLSI) guidelines (CLSI 2018). Most of the isolates were categorized as multidrug-resistant (resistant to three or more antimicrobial classes, Supplementary Table 1). As reference for differential gene expression and sequence variation analysis the UCBPP-PA14 strain was chosen.

### Colony screening

To rule out possible contaminations, all isolates were continuously re-streaked at least twice from single colonies. Only isolates with reproducible outcomes in phenotypic tests were included in the final panel, which furthermore passed DNA sequencing quality control (>85 % sequencing reads mapped to *P. aeruginosa* UCBPP-PA14 reference genome, total read GC content of 64-66 %).

### RNA sequencing

For comparable whole transcriptome sequencing, all clinical isolates and the UCBPP-PA14 reference strain were cultivated at 37 °C in LB broth and harvested in RNAprotect (Qiagen) at OD_600_= 2. Sequencing libraries were prepared using the ScriptSeq RNA-Seq Library Preparation Kit (Illumina) and short read data (single end, 50 bp) was generated on an Illumina HiSeq 2500 machine creating on average 3 million reads per sample.

The reads were mapped with Stampy (v1.0.23; (Lunter and Goodson 2011)) to the UCBPP-PA14 reference genome (NC_008463.1), which is available for download from the Pseudomonas Genome database (http://v2.pseudomonas.com). Mapping and calculation of reads per gene (rpg) values was performed as described previously (Khaledi et al. 2016). Expression counts were log-transformed (to deal with zero values we added one to the expression counts).

### DNA sequencing

Sequencing libraries were prepared from genomic DNA using the NEBNext Ultra DNA Library Prep Kit (New England Biolabs) and sequenced in paired end mode on Illumina HiSeq or MiSeq machines, generating either 2×250 or 2×300 bp reads. On average 2.89 million reads were generated per isolate (ranging from 653,062 to 21,086,866 reads with at least 30 times total genome coverage per isolate). All reads were adapter and quality clipped using fastq-mcf (Andrews 2010).

All sequencing data are available at NCBI’s Gene Expression Omnibus (GEO) and Sequence Read Archive (SRA) under the accession numbers GSE123544 (RNA-sequencing reads) and PRJNA526797 (DNA-sequencing reads), respectively.

### SNP calling

DNA-sequencing reads were mapped with *Stampy* as described above (see RNA-sequencing). For variant calling, SAMtools, v 0.1.19 (H. Li et al. 2009) was used. Heterozygous single-nucleotide variants were converted to the most likely homozygous state.

### Phylogeny

Paired-end reads (read length 150, fragment size 200) of eight reference genomes were simulated using art_illumina (v2.5.8) with the default error profile at 20-fold coverage (Huang et al. 2012). Together with our 414 clinical isolates, the sequencing reads were mapped to the coding regions of reference genome UCBPP-PA14 by BWA-MEM (v0.7.15) (H. Li 2013). SAMtools (v1.3.1) (H. Li et al. 2009) and BamTools (Barnett et al. 2011) (v2.3.0) were used for indexing and sorting the aligned reads respectively followed by variant calling using FreeBayes (v1.1.0) (Garrison and Marth 2012). The consensus coding sequences were computed by BCFtools (v1.6) (H. Li 2011) and then sorted into families by corresponding reference regions. A gene family was excluded if the gene sequence of any of its member differed by more than 10% in lengths as compared to the length of the reference genome gene family. Totally, 5,936 families were retained. The sequences of each family were aligned by MAFFT (v7.310) (Katoh and Standley 2013), and the alignments were concatenated. SNP sites that were only present in a single isolate were removed from the alignment. The final alignment was composed of 558,483 columns, and the approximately maximum likelihood phylogeny was then inferred by FastTree (v2.1.10, double precision) (Price, Dehal, and Arkin 2010).

### Pan-genome analysis and indel calling

The trimmed reads were assembled with SPAdes, v.3.0.1 using the --careful parameter (Bankevich et al. 2012). The assembled genomes were annotated using Prokka (v1.12) (Seemann 2014) using the metagenome mode of Prokka for gene calling, as we had noticed that genes on resistance cassettes were often missed by the standard isolate genome gene calling procedure. The gene sequences were clustered into gene families using Roary (Page et al. 2015). We observed that Roary frequently clustered together gene sequences of drastically varying lengths due to indels or start and stop codon mutations in those gene sequences and frequently also splits orthologous genes into more than one gene family. To overcome this behaviour we modified Roary to require at least 95% alignment coverage in the BLAST step (https://github.com/hzi-bifo/Roary).

For matching the Prokka annotation and the reference annotation of the *PA14* strain we used bedtools (Quinlan 2014) to search for exact overlaps of the gene coordinates. In a second step we identified all Roary gene families that contained a *PA14* gene. To identify insertions and deletions in the Roary gene families we extracted nucleotide sequences for each gene family and used MAFFT (Katoh and Standley 2013) to infer multiple sequence alignments. We restricted this analysis to gene families present in at least 50 strains. Then we used MSA2VCF (https://github.com/lindenb/jvarkit/) for calling variants in the gene sequences and restricted the output to insertion and deletions of at least nine nucleotides.

### Support vector machine classification

For applying cross-validation, the data set was split once randomly and once phylogenetically informed (see below) into k-folds (k set to 10, unless specified otherwise). Classifier hyperparameters were optimized on a k-1 fold sized partition, and performance of the optimally parameterized method was determined on the left out k fraction of the data. This was performed for all possible k partitions, assignments summarized and final performance measures obtained by averaging.

### Comparison of different machine learning classifiers

We used the training set for hyperparameter tuning of the classifiers, i.e. a linear SVM, RF, and LR, optimizing the F1-score in ten-fold cross-validation and then evaluated the best trained classifier on the held-out set. The RF classifier was optimized for the macro F1-score over different hyperparameters: (i) the number of decision trees in the ensemble, (ii) the number of features for computing the best node split, (iii) the function to measure the quality of a split and (iv) the minimum number of samples required to split a node. The logistic regression and the linear SVM were optimized for the macro F1-score over: (i) the C parameter (inverse to the regularization strength) and (ii) class weights (to be balanced based on class frequencies or to be uniform over all classes). Subsequently we measured the performance of the optimized classifiers over accordingly generated, held-out sets of samples).

In clinical practice *P. aeruginosa* strains isolated from patients are likely to include sequence types that are already part of our isolate collection. To obtain a more conservative estimate of the performance of the antimicrobial susceptibility prediction, we also validated the classifiers on a held-out dataset composed of entirely new sequence types and also selected the folds in cross-validation to be non-overlapping in terms of their sequence types (block cross-validation). For partitioning the isolate collection into sequence types we used spectral clustering over the phylogenetic similarity matrix (von Luxburg 2007). We obtained this matrix by applying a Gaussian kernel over the matrix of distances between isolates based on the branch lengths in the phylogenetic tree.

### Multi Locus Sequence Typing (MLST)

Consensus fastq files for each isolate were created with SAMtools to extract the seven *P. aeruginosa* relevant MLST gene sequences (*acsA, aroE, guaA, mutL, nuoD, ppsA, trpE*). Sequence type information was obtained from the Pseudomonas aeruginosa MLST Database (https://pubmlst.org/paeruginosa/) (Jolley and Maiden 2010).

### Implementation

We encapsulated the sequencing data processing routines in a stand-alone package named seq2geno2pheno https://github.com/hzi-bifo/seq2geno2pheno. The SVM classification was conducted with Model-T https://github.com/hzi-bifo/Model-T. which is built on scikit-learn (Pedregosa et al. 2011) and was already used as the prediction engine in our previous work on bacterial trait prediction (Weimann et al. 2016). seq2geno2pheno also implements a framework to use a more broader set of classifiers, which we used to compare different classification algorithms for drug resistance prediction. Finally, we created a repository that includes scripts to re-produce the figures and analyses presented in this paper using the aforementioned packages https://github.com/hzi-bifo/Fighting_PA_AMR_paper.

## Supporting information

Supplemental Table S2 Classifier Comparison

Supplemental Table S6 Validation Performance

Supplemental Table S4 SVM Per Feature Performance

Supplemental Table S5 Best Feature Sets

Supplemental Table S3 SVM Performance

Supplemental Table S1 Isolate Strain Infos

## Author contributions

AxKo, MH, PG and GC generated data. AKh and MS performed experiments. AW, EA, MM and ACM developed the computational methodology. AW, AKh, MS, EA, TK, AB and MM and analysed the data. AW, AKh, MS, AO, ACM and SH interpreted the results. ACM and SH conceived the project, designed experiments and supervised the work. AW, AKh and TK generated figures and tables. AKh, AW, ACM and SH wrote the paper. All authors read and approved the final manuscript.

## Acknowledgements

Financial support from the European Research Council (http://erc.europa.eu/) (ERC COMBAT grant 724290) is gratefully acknowledged. This study was further supported by the German Federal Ministry of Education and Research (grant 01 KI 9907). Members of the study group on “Spread of nosocomial infections and resistant pathogens” contributed bacterial isolates. GC and AO are supported by Instituto de Salud Carlos III, Ministerio de Economía, Industria y Competitividad, Spanish Network for Research in Infectious Diseases (REIPI RD16/0016) and grant PI18/00076. We thank Adrian Kordes for his assistance in preparing the DNA sequencing libraries, Sarah Pohl for critical input on data processing and Agnes Nielsen for conducting the AST testing of the clinical isolates.

## Supplementary Information

**Supplementary Table S1:** Pseudomonas isolate resistance and supplier information. Isolate: the isolate name, Supplier (Geographic origin): Institution that provide each strain and the sampling site.

**Supplementary Table S2:** Machine classification performance across drugs, combination of data types and different cross-validation schemes (randomly: sheet random) or phylogenetically informed: sheet tree) for the training and validation data set (including a summary for the best data type combination for each drug (best data type combinations). Drug: one of four anti-pseudomonal drugs, Classifier: one of LR (logistic regression), SVM (support vector machine), RF (random forest), Data type: data type combination used, Macro F1 cv: macro F1-score based on cross-validation, macro F1 validation: macro F1-score based on the validation set.

**Supplementary Table S3:** Support vector machine classification performance across drugs, combination of data types and different cross-validation schemes for the training data set. cv_mode: cross-validation scheme, data type: data type combination, drug: anti-pseudomonal drug, measure: performance measure, value: performance measure value, std: standard deviation of performance measure.

**Supplementary Table S4:** Number of markers for different values of the SVM C-parameter for each data type combination. data type: data type combination, drug: anti-pseudomonal drug. no_feats: number of non-zero features in the model, F1-score_macro: overall performance measure, c_param: value of the SVM C parameter.

**Supplementary Table S5:** Genetic and expression markers for each data type combination. cparam: optimal SVM C parameter, data type: data type combination, drug: anti-pseudomonal drug, feature: marker id, which is a combination of PA14 reference gene id and gene name, Prokka/Roary gene id or name, SNP position and nucleotide and amino acid change (if applicable) and number of markers with identical distribution in the data set. SVM weight: quantitative contribution of the marker in the SVM model.

**Supplementary Table S6:** SVM classification performance across drugs, combination of data types for different performance measures for the validation data set. data type: data type combination, drug: anti-pseudomonal drug, c_param: the C parameter of the optimal SVM model used for prediction.

**Supplementary Figure 1:**
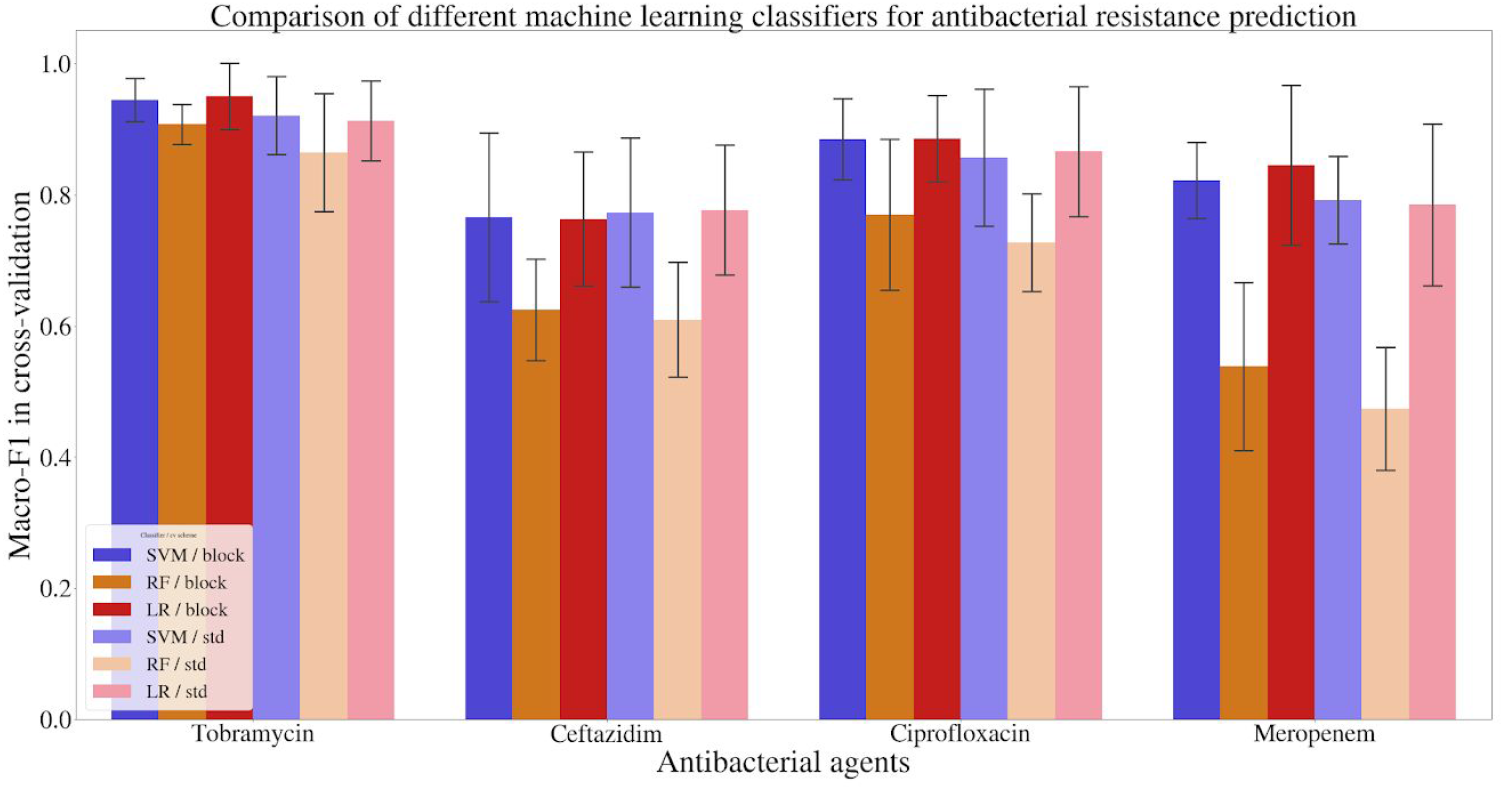
Comparison of support vector machine, random forest, and logistic regression classifiers in antimicrobial resistance prediction, and their generalization ability. Training and performance estimates was performed in ten-fold cross-validation, where isolates were split either randomly (standard: std) or using phylogenetically related blocks of isolates (block). The classification was performed for each antibiotic using the best performing feature combinations for the SVM. The error bar shows the variability (standard deviation) in 10 test folds.

**Supplementary Figure 2:**
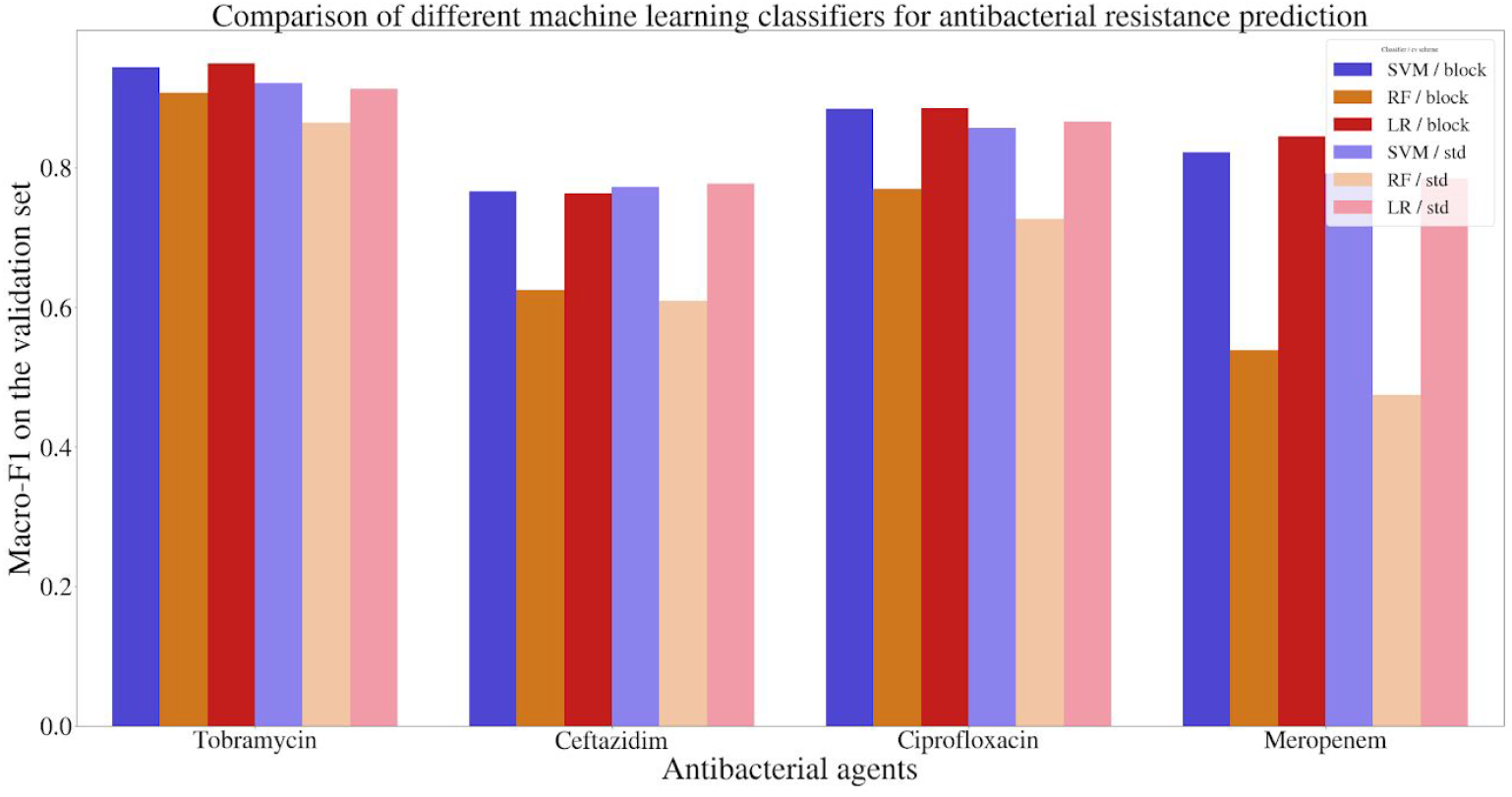
Comparison of support vector machine, random forest, and logistic regression classifiers in antimicrobial resistance prediction, and their generalization ability. The classifiers were tuned in a ten-fold cross-validation; subsequently their performances are reported over the held-out set, where isolates were split either randomly (standard: std) or using phylogenetically related blocks of isolates (block).

**Supplementary Figures S3-S6:**
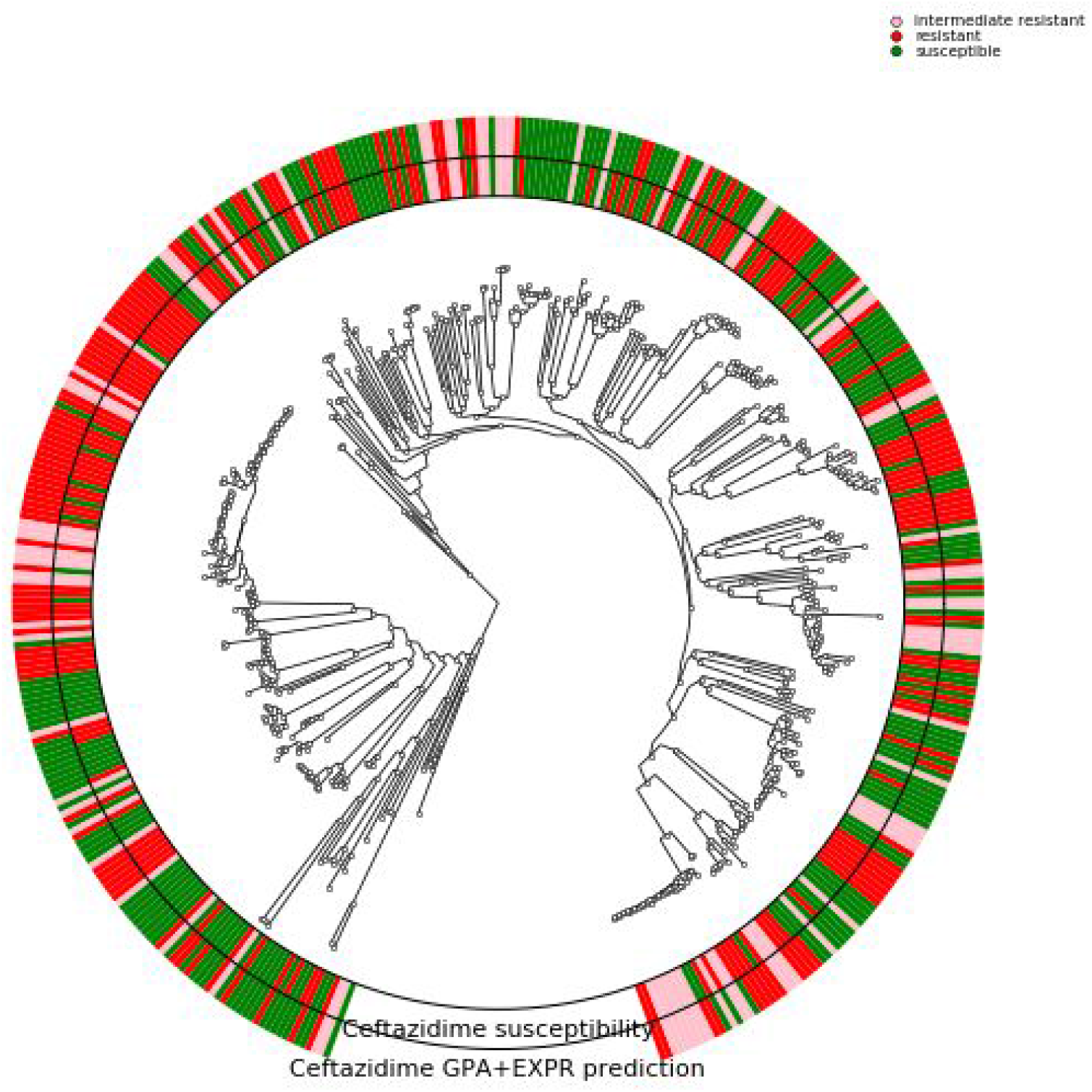
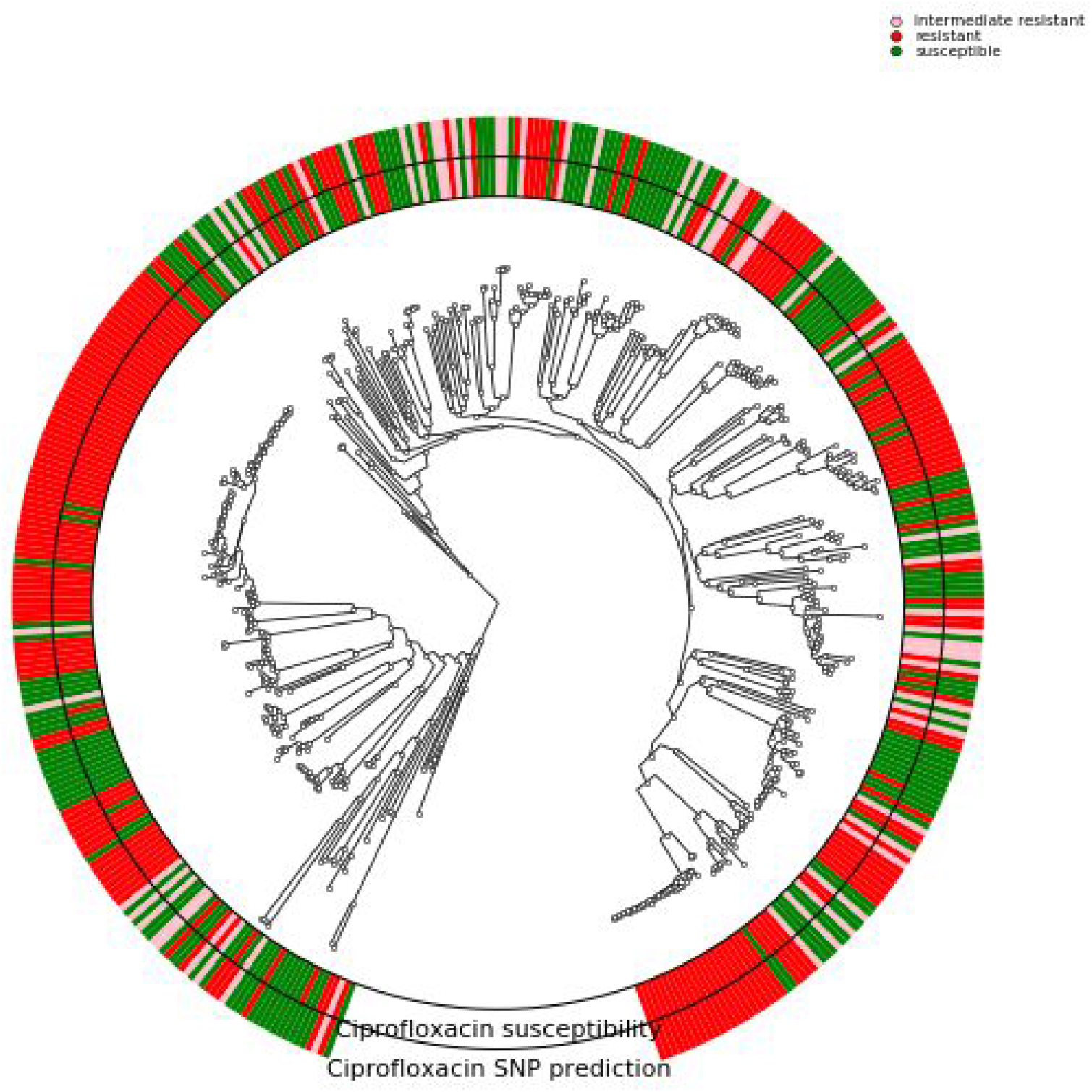
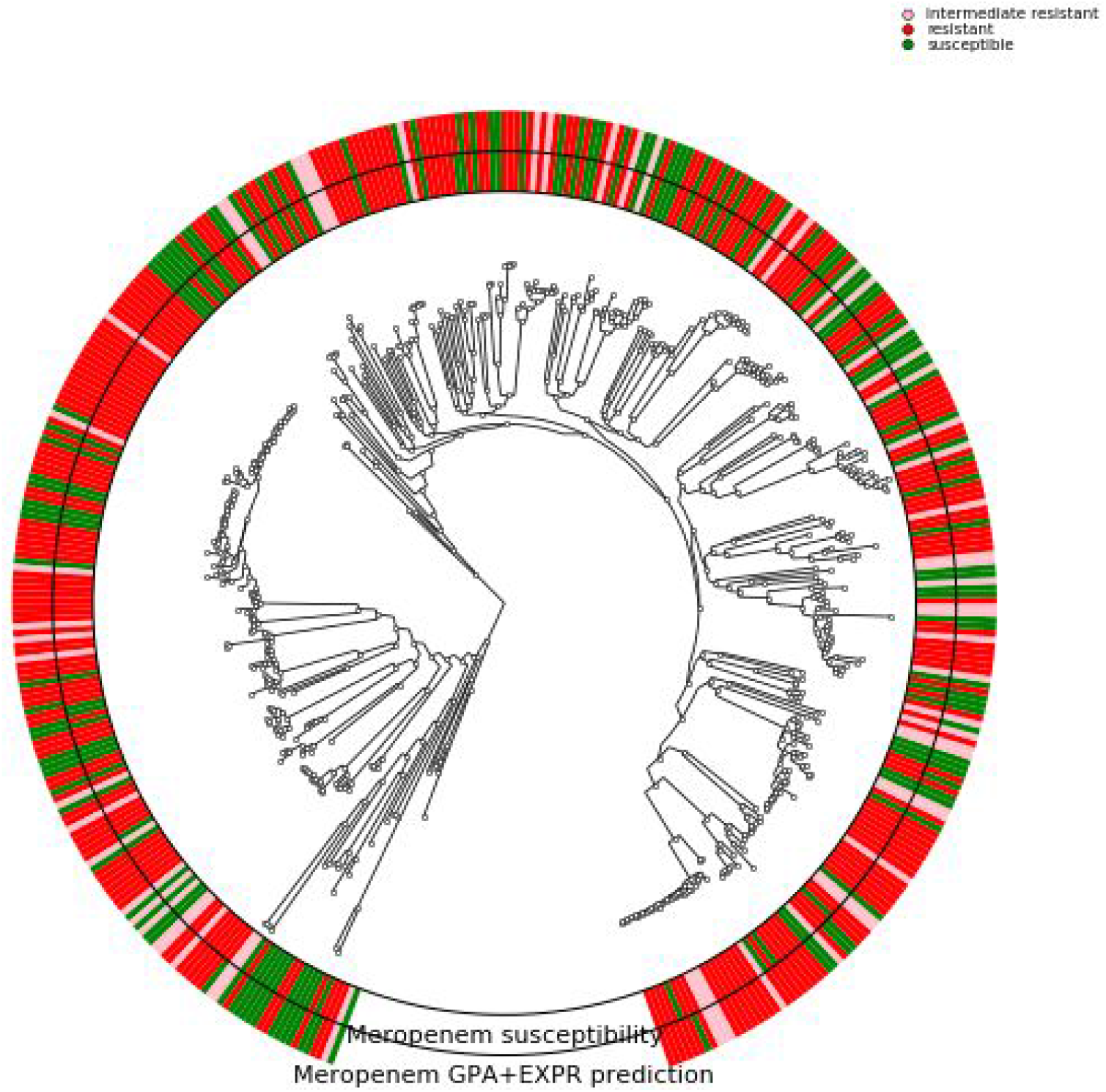
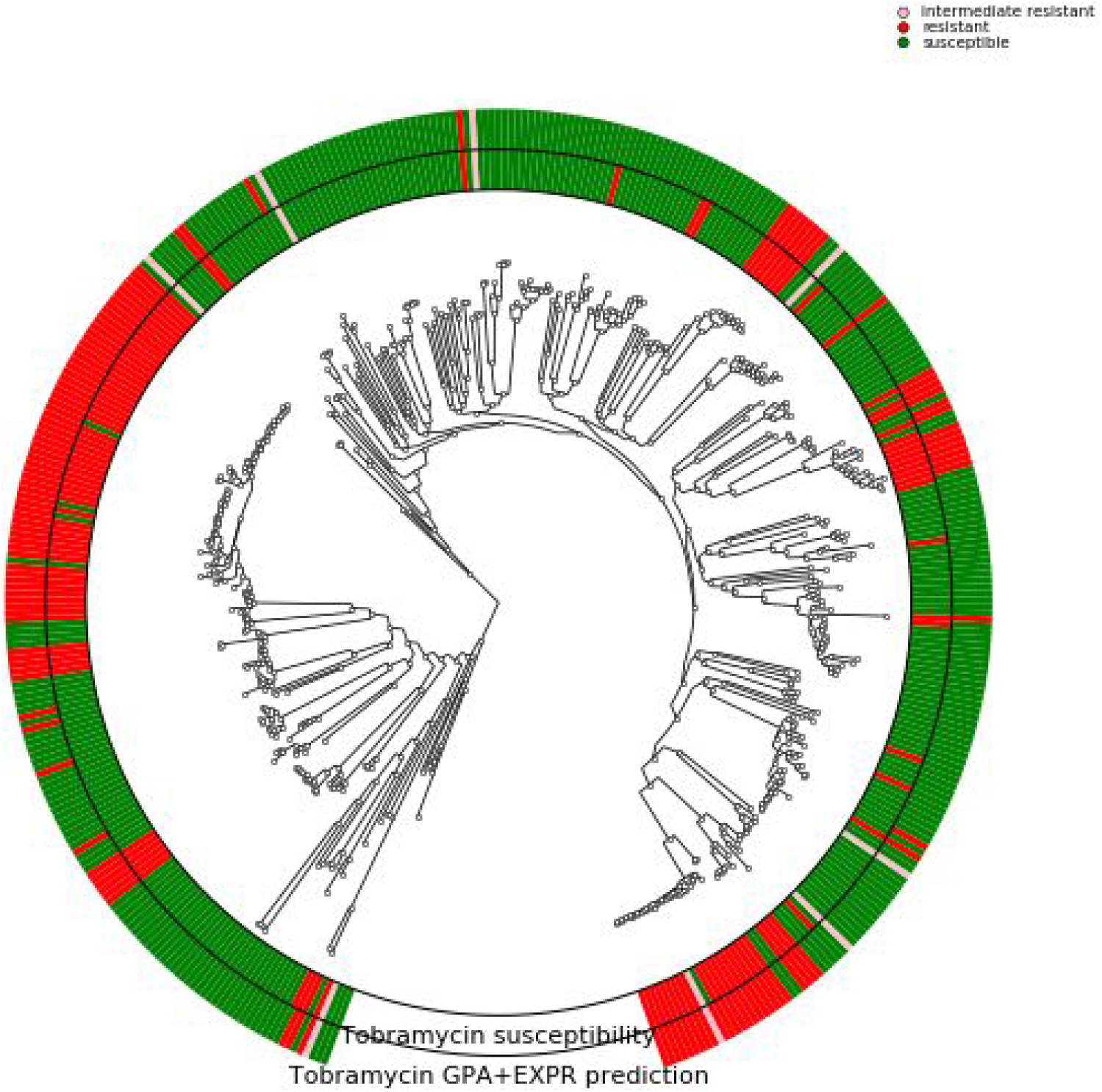
Phylogenetic tree of the 414 *P. aeruginosa* isolates used in this study. The branches leading towards two deeply branching clades were collapsed (CH4433 and ESP077, and CH4684, CH5206 and CH5387). The inner ring depicts the susceptibility of each isolate to ciprofloxacin, meropenem, tobramycin, and ceftazidime respectively; green susceptible, red resistant; rose intermediate resistant. The outer rings shows the susceptibility as assigned by the diagnostic genomic classifier (intermediate resistant samples were not assigned).

